# Renoprotective effects Of Dexmedetomidine against ischemia-reperfusion injury in streptozotocin-induced diabetic rats

**DOI:** 10.1101/325894

**Authors:** Seung Hyun Kim, Ji Hae Jun, Ju Eun Oh, Eun-Jung Shin, Young Jun Oh, Yong Seon Choi

## Abstract

**Background:** Diabetic patients are susceptible to renal ischemia-reperfusion injury, which leads to perioperative complications. Nucleotide binding and oligomerization domain (NOD)-like receptor 3 inflammasome participates in the development of diabetes, and contributes to renal ischemia-reperfusion injury. Dexmedetomidine, a highly selective α2-adrenoreceptor agonist, shows renoprotective effects against ischemia-reperfusion injury. We aimed to elucidate the effects, underlying mechanisms, and optimal timing of dexmedetomidine treatment in diabetic rats.

**Methods:** Male Sprague-Dawley rats (60 animals, weighing 250-300 g) were randomly divided into normal-sham, diabetes-sham, diabetes-ischemia-reperfusion-control, diabetes-ischemia-reperfusion-dexmedetomidine-pre-treatment, and diabetes-ischemia-reperfusion-dexmedetomidine-post-treatment groups. Renal ischemia-reperfusion injury was induced in diabetic rats by occlusion of both renal arteries for 45 minutes followed by reperfusion for 24 hours. Dexmedetomidine (10 μg/kg) was administered intraperitoneally 1 hour before ischemia (pre-treatment) or upon reperfusion (post-treatment). After reperfusion, renal tissue was biochemically and histopathologically evaluated.

**Results:** Dexmedetomidine treatment attenuated IR-induced increase in NLRP3, caspase-1, IL-1β, phospho-AKT, and phospho-ERK signaling. Moreover, oxidative stress injury, inflammatory reactions, apoptosis, and renal tubular damage were favorably modulated by dexmedetomidine treatment. Furthermore, post-reperfusion treatment with dexmedetomidine was significantly more effective than pre-treatment in modulating inflammasome, AKT and ERK signaling, and oxidative stress.

**Conclusions:** This study shows that protective effects of dexmedetomidine in renal ischemia-reperfusion injury are preserved in diabetic conditions and may potentially provide a basis for the use of dexmedetomidine in clinical treatment of renal ischemia-reperfusion injury.

## Introduction

Acute kidney injury (AKI) is a common complication in surgical patients and is associated with increased mortality, risk of chronic kidney disease and hemodialysis after discharge, and cost. Diabetes mellitus is a metabolic disorder with major complications including microvascular and macrovascular diseases and an important risk factor for AKI resulting in renal damage and dysfunction, as it increases susceptibility to ischemia-reperfusion (IR) injury.[1]

Diabetic nephropathy is the leading cause of end-stage renal disease. In renal tissues of streptozotocin (STZ)-induced diabetic rats, nucleotide binding and oligomerization domain (NOD)-like receptor 3 inflammasome is activated. Suppression of NLRP3 inflammasome activation was shown to significantly reduce renal tissue inflammation and improve renal function.[2] NLRP3 knock-out mice were protected from renal tubule damage and renal interstitial inflammation in kidney unilateral ureteral occlusion and IR models.[3] These data show that NLRP3 inflammasome plays a key role in kidney IR injury.

Dexmedetomidine (DEX) is a highly selective α2-adrenoreceptor agonist with sedative, analgesic, sympatholytic, and hemodynamic stabilizing properties. Previous studies suggest that DEX has organoprotective effects, reducing cerebral and cardiac injury.[4,5] DEX also reduced renal injury, an effect abolished by α2-adrenoreceptor antagonist atipamezole.[6] Moreover, DEX protected against kidney IR injury through anti-apoptotic, anti-inflammatory, and antioxidant effects in diabetic rats.[7] Despite clinical importance of mitigating renal damage in IR injury in surgical patients, underlying molecular mechanisms of DEX remain poorly understood.

Here, we investigate the protective effects of DEX against renal IR injury in diabetic rats and elucidate the relation between DEX and NLRP3 inflammasome. Additionally, this study attempted to identify DEX treatment regimens achieving optimal renoprotective effects in a diabetic rat renal IR injury model.

## Materials and methods

### Animal preparation

All animal experiments were approved by the Committee for the Care and Use of Laboratory Animals, Yonsei University College of Medicine (No. 2015-0129, approval date 16.06.2015), and were performed in accordance with the *Guide for the Care and Use of Laboratory Animals* published by the US National Institutes of Health.[8]

Male Sprague-Dawley rats (60 animals, 10-12 weeks old, 250-300 g) were used. Diabetes was induced by a single intraperitoneal injection of freshly prepared STZ (85 mg/kg) dissolved in 100 mM sodium citrate buffer (pH 4.5), and operations were performed 2 weeks after STZ administration. Blood sugar concentration was determined using tail blood samples. Rats were housed in a temperature and light-controlled animal facility with alternating 12-hour light/dark cycles and *ad libitum* access to standard laboratory chow and water.

General anesthesia was induced using xylazine (Rompun, 10 mg/kg) and tiletamine + zolazepam (Zoletil 50, Virbac Korea, 30 mg/kg). Rats were intubated with a 16-gauge (G) catheter and artificially ventilated (Harvard Apparatus 683, Holliston, MA, USA) at 30-35 cycles/min. The right femoral artery was cannulated to monitor mean arterial pressure (MAP) and collect blood. Heart rate (HR) was monitored by subcutaneous stainless-steel electrodes connected to the power lab system (ML845 PowerLab with ML132; AD Instruments, Colorado Springs, CO, USA). Body temperature was maintained at approximately 37 °C using a heating pad and continuously monitored throughout the experiment.

### Experimental models and study groups

Rats were randomly divided (12 rats per group) into normal-sham (N-sham), diabetic-sham (D-sham), diabetic IR control (D-IRC) receiving normal saline, diabetic DEX pre-treatment-IR (D-DEXpre), and diabetic IR-DEX post-treatment (D-DEXpost) groups. We used a previously described rodent model of renal IR injury.[9] In brief, both rat renal arteries were clamped for 45 min with non-traumatic microvascular clamps followed by 24 h of reperfusion. Ischemia and reperfusion were confirmed by visually inspecting the kidneys. MAP and HR were continuously monitored and recorded during the procedures (baseline, during ischemia, and after reperfusion). All diabetic rats underwent midline incision to expose both kidneys. To examine DEX renoprotective effects, DEX (Precedex, Pfizer, 10 μg/kg) was administered 1 h before ischemia or upon reperfusion, whereas the control group received an equivalent amount of normal saline. Initially, we tested the efficacy of various DEX doses (10, 20, 50, and 100 μg/kg) 1 h before ischemia and found that 10 μg/kg yielded optimal renoprotective effects against renal tubular damage. IR-control groups received equivalent amounts of normal saline via tail vein. Blood glucose concentration >200 mg/dl was considered high glucose level.[10] Blood glucose concentrations were determined at baseline, before ischemia, upon reperfusion, and 24 h later. Kidneys were collected after 24-h reperfusion.

### Periodic acid-Schiff staining

Tissue samples for histopathological examination were taken from the left kidney (including the ischemic zone) after 24-h reperfusion, fixed in 10% buffered formalin, and embedded in paraffin. Paraffin-embedded kidney tissues were cross-sectioned through the midpoint to analyze histologic damage. Periodic acid-Schiff (PAS) staining was performed as previously described.[11] Degree of tubular damage was assessed on a scale of 1 to 4 (no histopathological findings observed, focal area of tubular necrosis involving <25% of the kidney, tubular necrosis involving 25-50% of the kidney, and tubular necrosis involving >50% of the kidney corresponding to 1-4, respectively)[11] (N=6).

### TUNEL assay

Apoptosis was detected in paraffin sections from each group using the terminal deoxynucleotidyl transferase (TdT)-mediated uridine triphosphate (dUTP) nick end labeling (TUNEL) as previously described.[12] Sections (5 μm) were stained using in situ DeadEnd™ Colorimetric Apoptosis Detection System (Promega, Madison, WI, USA) according to the manufacturer’s instructions. Five visual fields from each sample block were randomly selected and analyzed by a blinded observer using an Olympus microscope with 400× magnification. Apoptotic index, defined as (apoptotic cells/total cells) × 100%, was determined using 20 fields per sample (N=6).

### BUN and creatinine analysis

BUN and serum creatinine concentrations were determined 24 h after reperfusion using picric acid and diacetyl monoxime methods[13], respectively.

### Immunoblotting

Tissue samples were lysed in RIPA buffer (10 mM Tris-HCl (pH 7.5), 1 mM EDTA, 150 mM NaCl, 1% NP-40, 2% SDS, 1% sodium deoxycholate, 50 mM NaF, 0.2 mM Na_3_VO_4_, 1 mM PMSF, and phosphatase inhibitor cocktail I and II (Sigma, St. Louis, MO, USA). Protein amounts were determined using the Quick Start Bradford 1× Dye reagent (Bio-Rad, USA).

Proteins were separated on sodium dodecyl sulfate-polyacrylamide gel electrophoresis (SDS-PAGE) and immunoblotted with anti-phospho-eNOS, anti-eNOS, anti-phospho-iNOS, anti-iNOS, anti-Bcl-2, anti-Bax, anti-PARP, anti-cleaved-PARP, anti-interleukin 1β, anti-phospho-AKT, anti-AKT, anti-phospho-ERK, anti-ERK, and anti-actin antibodies obtained from Cell Signaling Technology (Beverly, MA, USA) and anti-CTP1A, anti-NOX4, anti-TXNIP, anti-NLRP3, anti-ASC, anti-caspase-1, and anti-cleaved caspase-1 antibodies obtained from Abcam (Cambridge, UK) (N = 6).

### ELISA

Serum levels of interleukin (IL)-6 and tumor necrosis factor (TNF)-α were determined by commercial ELISA kits (R&D Systems, Minneapolis, MN, USA) according to the manufacturer’s instructions.

### Statistical analysis

Data are expressed as means ± S.D. and were analyzed using one-way analysis of variance (ANOVA) followed by Bonferroni correction or repeated measures two-way ANOVA. P-values <0.05 were considered significant.

## Results

### Hemodynamic parameters

Diabetic rats displayed increased blood glucose levels and reduced body weight. Table 1 shows hemodynamic data for all groups throughout the study period. No significant differences in MAP and HR at baseline or during ischemia were observed between groups. MAP was increased by IR; however, post-reperfusion treatment with DEX significantly attenuated this increase (p <0.05). IR-induced HR increase was significantly attenuated in both DEX-treated groups (p <0.05).

**Table 1.** Changes in hemodynamic parameters (means ± SD) in diabetic rats. MAP and HR were recorded 15 min after starting intraperitoneal anesthesia (baseline), 45 min after inducing ischemia (during ischemia), and after 24-h reperfusion (after reperfusion). MAP: mean arterial blood pressure; HR: heart rate; DEX: dexmedetomidine; D-IRC: diabetic ischemia-reperfusion control. *p <0.05, compared with the sham baseline; †p <0.05, compared with D-IRC group; ‡p <0.05, compared with DEX pre-treatment group

| MAP | D-Sham | D-IRC | DEX<sup>pre</sup> | DEX<sup>post</sup> |
| --- | --- | --- | --- | --- |
| Baseline | 90.2 ± 17.7 | 89.4 ± 9.6 | 96.3 ± 12.2 | 90.7 ± 7.0 |
| During ischemia | | 84.7 ± 23.1 | 81.1 ± 5.1 | 73.9 ± 11.0 |
| After reperfusion | | 120.6 ± 7.4<sup>*</sup> | 123.2 ± 8.1<sup>*</sup> | 114.1 ± 4.4<sup>*,‡</sup> |

| HR | D-Sham | D-IRC | DEX<sup>pre</sup> | DEX<sup>post</sup> |
| --- | --- | --- | --- | --- |
| Baseline | 269.3 ± 20.8 | 281 ± 25.9 | 314.5 ± 99.7 | 265.4 ± 23.8 |
| During ischemia | | 254.1 ± 71.7 | 230.0 ± 49.7 | 238.5 ± 29.8 |
| After reperfusion | | 312.1 ± 26.8<sup>*</sup> | 263.6 ± 30.5<sup>†</sup> | 257.9 ± 27.1<sup>†</sup> |

### DEX reduced renal tubular apoptosis in diabetic rats

No apoptotic cells were observed in kidney tissues of the normal sham-operated group. A small number of apoptotic cells were observed in the diabetic sham-operated group; however, the difference between the sham groups was not statistically significant. IR injury significantly increased the number of apoptotic tubular cells in diabetic rats. High-dose DEX treatments (20, 50, and 100 μg/kg) did not reduce apoptosis (Figure 1A), whereas 10 μg/kg of DEX significantly ameliorated IR-induced apoptosis (p <0.05). DEX post-reperfusion treatment group displayed significantly reduced tubular cells apoptosis, compared to DEX pre-treatment group (p <0.05, Figure 1B).

**Figure 1.**
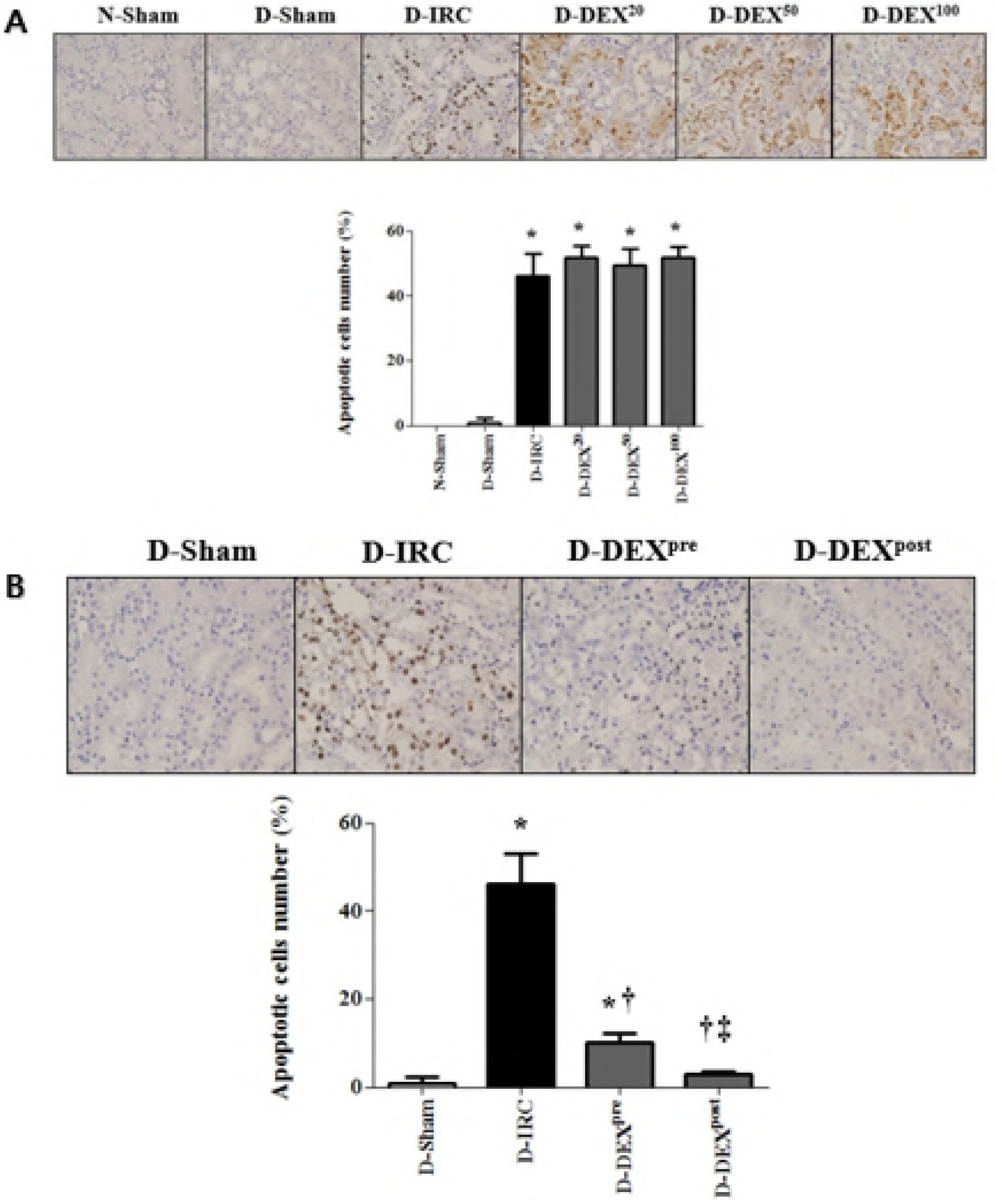
DEX reduced IR-induced renal tubular cell death in diabetic rats, TUNEL X40. High-dose DEX treatments (20, 50, and 100 μg/kg) did not reduce apoptosis (A). DEX (10 μg/kg) reduced apoptosis (B). DEX^20^, DEX^50^, and DEX^100^ correspond to DEX treatments of 20, 50, and 100 μg/kg, respectively; *p <0.05, compared with D-sham baseline; †p <0.05, compared with D-IRC group; ‡p <0.05, compared with DEX pre-treatment group

### DEX improved renal tubular damage following IR in diabetic rats

Histopathological scores of renal tubular damage were significantly higher in D-IRC group, compared to D-sham group scores (Figure 2A). Post-reperfusion DEX treatment significantly improved tubular damage compared to damage in the D-IRC group (p <0.05).

### Post-reperfusion treatment with DEX attenuated IR-induced renal dysfunction in diabetic rats

BUN and serum creatinine levels observed 24 h after IR were significantly higher than those measured in D-sham group (p <0.05). Compared to untreated D-IRC group, post-reperfusion treatment with DEX significantly reduced BUN and creatinine levels (p <0.05) (Figure 2B).

### DEX reduced serum levels of IL-6 and TNF-α following IR injury in diabetic rats

Following IR, serum IL-6 and TNF-α levels were significantly increased in D-IRC group compared with D-sham group (p <0.05). Pre- and post-reperfusion DEX treatments significantly decreased IR-induced elevation of IL-6 and TNF-α (p <0.05) (Figure 2C).

**Figure 2.**
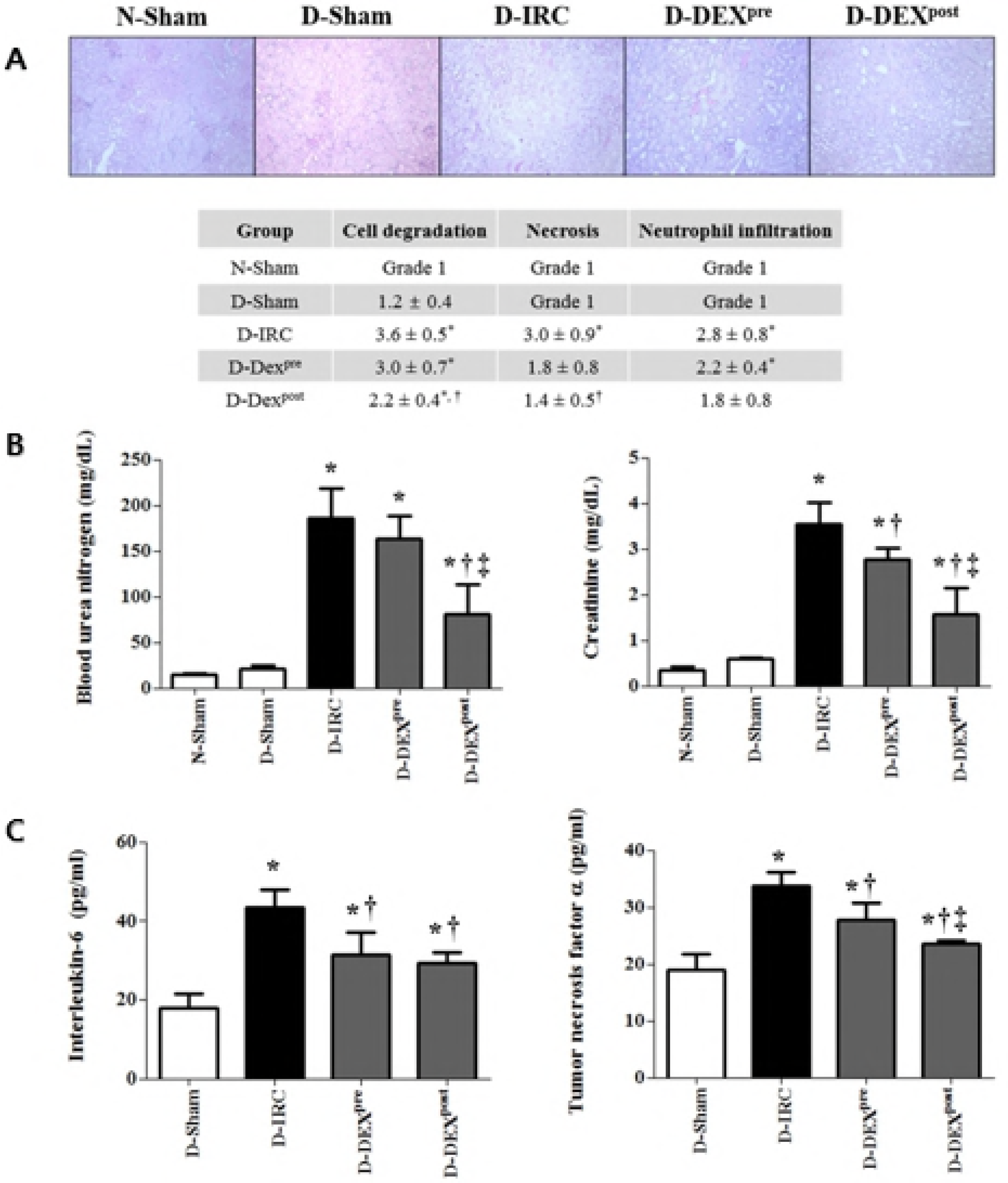
DEX improved histopathological scores and reduced markers of renal dysfunction and inflammation in renal IR injury, PAS X40. Post-reperfusion treatment with DEX improved histopathological scores of renal tubular damage in diabetic rats (A). DEX reduced BUN and creatinine (B) and IL-6 and TNF-α levels (C) in renal IR injury. *p <0.05, compared with D-sham baseline; †p <0.05, compared with D-IRC group; ‡p <0.05, compared with DEX pre-treatment group

### DEX did not affect expression of p-eNOS and attenuated expression of p-iNOS in diabetic rats

Levels of p-eNOS were decreased in the D-IRC group compared to p-eNOS levels of the D-sham group, whereas p-eNOS expression in both pre-treatment and post-reperfusion DEX treatment groups was retained. Expression of p-iNOS was barely detectable in the D-sham group and significantly upregulated in the D-IRC group (p <0.05). DEX pre-treatment and post-reperfusion treatment both significantly decreased p-iNOS expression compared to expression in the D-IRC group (p <0.05, Figure 3A).

### DEX favorably altered expression of Bcl-2, Bax, and cleaved-PARP in diabetic rats

Compared with D-sham group, Bcl-2 expression was significantly reduced in D-IRC group, and significantly higher in DEX post-reperfusion treatment group than in D-IRC group (p <0.05). In contrast, cleaved-PARP and PARP expression was significantly increased in D-IRC group compared with D-sham group, effect ameliorated by DEX post-reperfusion treatment (p <0.05) (Figure 3B). Both pre- and post-reperfusion treatment with DEX ameliorated IR-induced increase of Bax expression, compared to D-IRC group (p <0.05), with post-reperfusion DEX treatment proving more effective (p <0.05).

**Figure 3.**
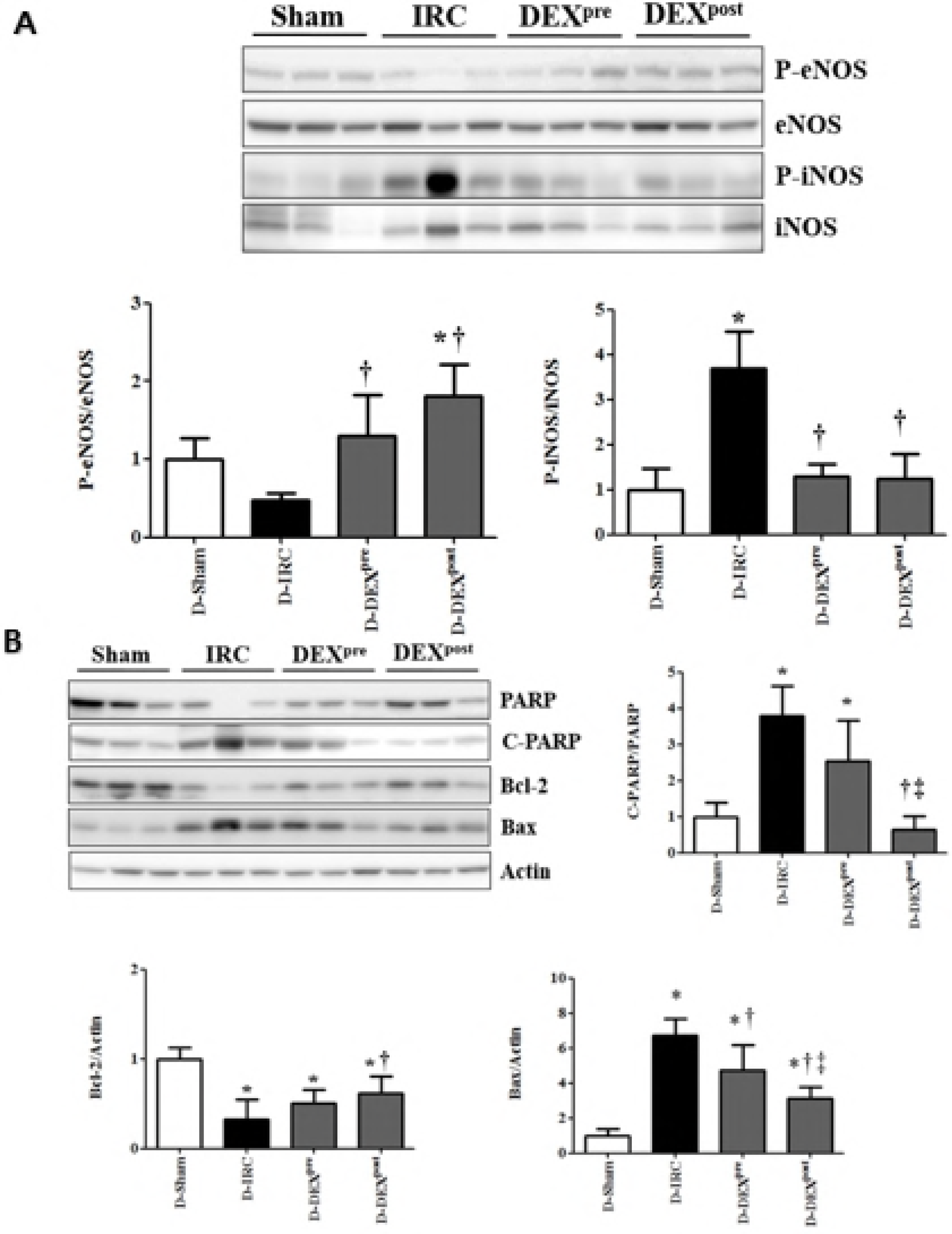
Effects of DEX on NOS, cleaved-PARP, Bcl-2, and Bax in renal IR injury. DEX did not affect expression of p-eNOS and attenuated expression of p-iNOS in diabetic rats (A). Post-reperfusion treatment with DEX reduced the increase in cleaved-PARP and did not affect Bcl-2 and Bax expression in renal IR injury (B). *p <0.05, compared with D-sham baseline; †p <0.05, compared with D-IRC group; ‡p <0.05, compared with DEX pretreatment group

### Post-reperfusion DEX treatment attenuated increase in CTP1A, NOX4, and TXNIP levels following IR in diabetic rats

Following IR, CTP1A, NOX4, and TXNIP levels were significantly increased in D-IRC group compared with D-sham group, an effect significantly reduced in DEX post-reperfusion treatment group (p <0.05, Figure 4A).

### Post-reperfusion DEX treatment decreased IR-induced increase in phospho-AKT and phospho-ERK in diabetic rats

Pre- and post-reperfusion DEX treatments ameliorated IR-induced increase in phospho-AKT and phospho-ERK compared to D-IRC group levels (p <0.05), with post-reperfusion treatment with DEX proving more effective (p <0.05, Figure 4B).

**Figure 4.**
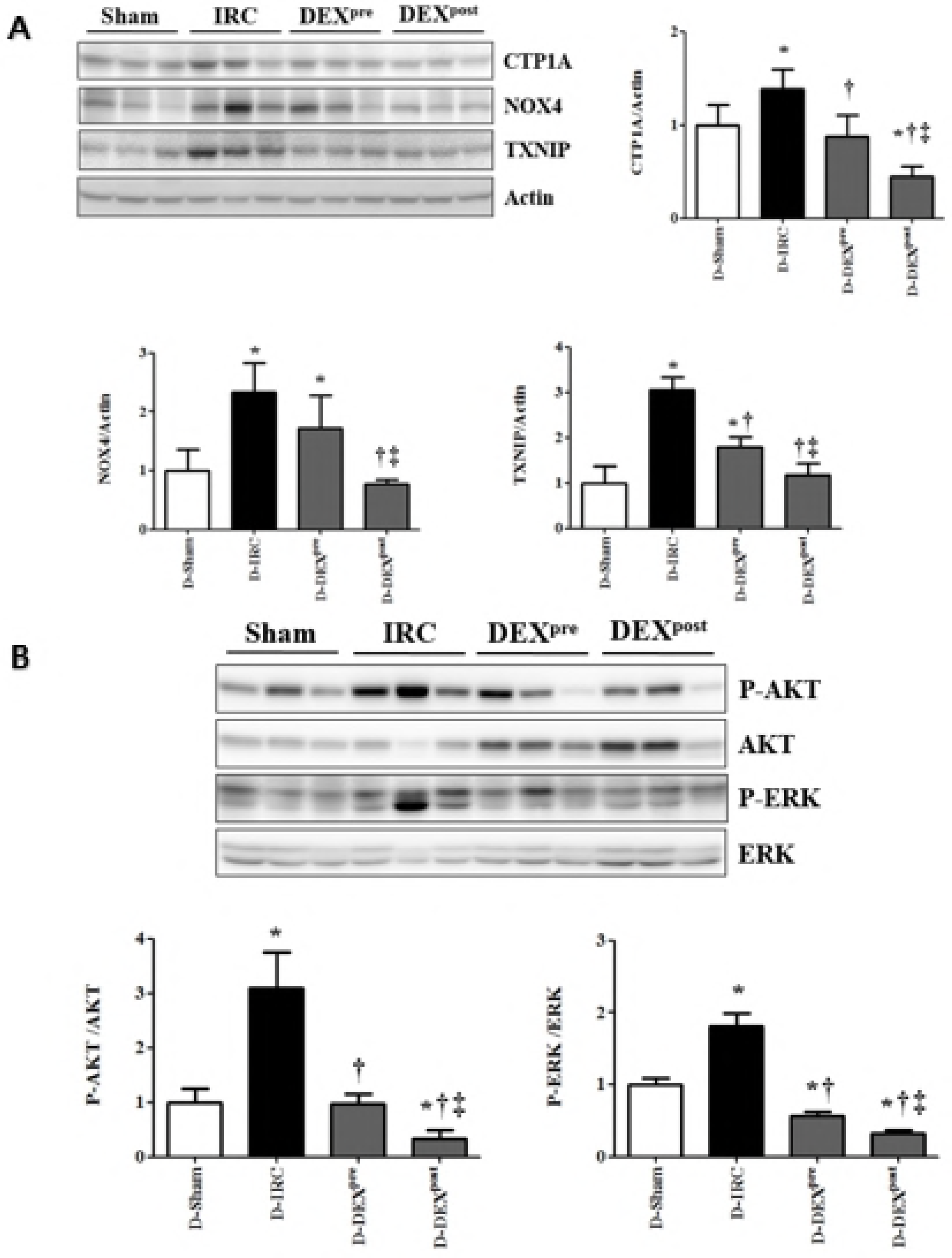
Effects of DEX on oxidative stress markers, AKT and ERK signaling in renal IR injury. Post-reperfusion DEX treatment attenuated the increase in CTP1A, NOX4, and TXNIP in renal IR injury (A) and reversed the increase in phospho-AKT and phospho-ERK levels in renal IR injury (B). *p <0.05, compared with D-sham baseline; †p <0.05, compared with D-IRC group; ‡p <0.05, compared with DEX pre-treatment group

### Post-reperfusion DEX treatment ameliorated increase in NLRP3, cleaved caspase-1, and IL-1β levels following IR in diabetic rats

IR-induced increase in levels of NLRP3, cleaved caspase-1, and IL-1β was significantly diminished in DEX post-reperfusion treatment group compared with D-IRC group (p <0.05). No significant difference in ASC levels was observed between diabetic rat groups (Figure 5).

**Figure 5.**
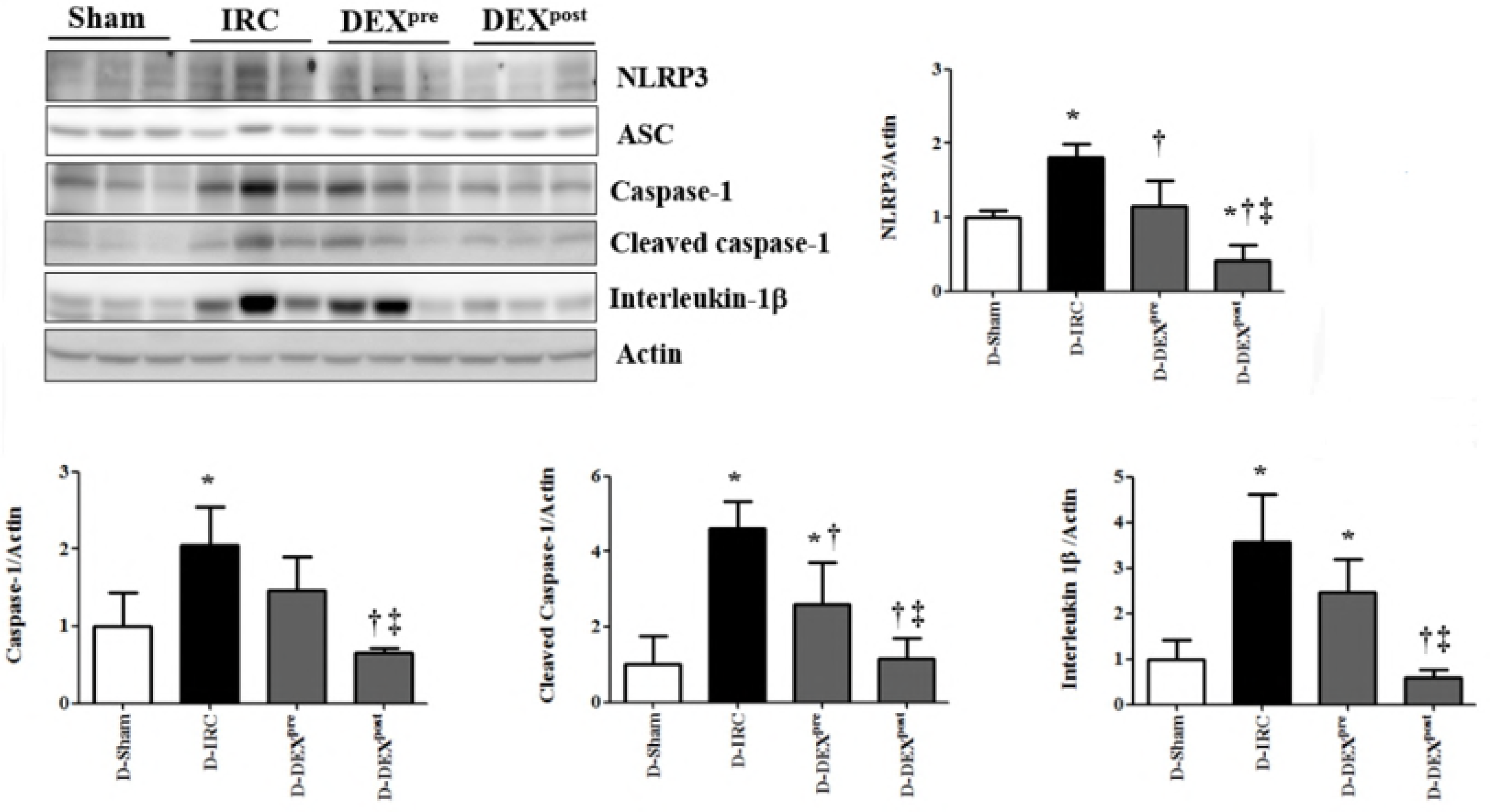
Post-reperfusion DEX treatment ameliorated the increase in NLRP3, cleaved caspase-1, and IL-1β in renal IR injury. *p <0.05, compared with D-sham baseline; †p <0.05, compared with D-IRC group; ‡p <0.05, compared with DEX pre-treatment group

## Discussion

This is the first study to demonstrate that renoprotective effects of DEX in renal IR injury are associated with the inflammasome. IR injury increased the levels of inflammasome components NLRP3 and caspase-1, an effect reversible by DEX. DEX also reduced increased IL-1β levels, activated by the inflammasome. Inflammasome-associated protective effect of DEX was greater in the post-reperfusion treatment group than in the pre-treatment group. Post-reperfusion DEX treatment had greater protective effect than pre-treatment in terms of AKT and ERK signaling, oxidative stress, and renal dysfunction.

NLRP3 inflammasome contributes to the development of type 1 and 2 diabetes mellitus.[14,15] The inflammasome is a promising potential therapeutic target in numerous renal diseases. NLRP3 protein, the essential component of the inflammasome, contributes to renal IR injury via direct effects on renal tubular epithelium.[3] Chen et al. analyzed renal interstitial inflammation in diabetes patients and found increased levels of NLRP3, IL-1β, and IL-18 in diabetic nephropathy, compared to normal controls.[2] Inhibition of NLRP3 activation protects diabetic kidney from IR injury and diminishes sensitivity of diabetic kidney tissues to AKI.[16] Gu et al. have demonstrated that DEX provided cytoprotection and improved tubular architecture and function following renal ischemia.[6] Thus, we hypothesized that inhibition of inflammasome might retain DEX protective effect against IR injury in STZ-induced diabetic rat models.

The inflammasome is a complex of cytosolic proteins consisting of an NLR (Nod-like receptor), an adaptor protein (ASC), and caspase-1. When NLRP3 inflammasome is activated, caspase-1 and subsequently IL-1β are cleaved to their active forms, stimulating inflammatory cascades. Here, IR injury increased NLRP3, caspase-1, cleaved caspase-1, and IL-1β levels, whereas inflammasome activation was suppressed by DEX treatment, indicating that protective effects of DEX are well preserved under diabetic conditions. Although DEX renoprotective effects against IR injury in diabetic rats are achieved by ameliorating lipid peroxidation and oxidative stress[7], present study is the first to demonstrate that protective effects of DEX against renal IR injury in diabetic rats are associated with the inhibition of inflammasome. This result is consistent with recent studies with non-diabetic animal models that demonstrated that DEX reduces inflammasome activation.[17]

DEX ameliorated IR-induced increase in phospho-AKT and phospho-ERK compared to D-IRC group levels and IR-induced decrease in anti-apoptotic Bcl-2 and pro-apoptotic Bax levels. Increase in cleaved-PARP was reduced by DEX post-reperfusion treatment. Reperfusion injury salvage kinase (RISK) pathway, combining two parallel cascades, phosphatidylinositol-3 kinase (PI3K)-AKT and p42/p44 extracellular signal-regulated kinase (ERK), is an important pro-survival pathway in IR injury. However, AKT activity increases in diabetes.[18] Kim et al. demonstrated that increased p-AKT in diabetes is accompanied by decreased anti-apoptotic Bcl-2 and pro-apoptotic Bax levels.[19] Increased ERK activity was observed in human and rodent diabetic adipose tissue, and inhibition of the ERK pathway is a potential therapeutic strategy to combat insulin resistance.[20] Reactive oxygen species (ROS) produced during IR injury trigger over-activation of the nuclear enzyme poly (ADP-ribose) polymerase (PARP), which induces translocation of apoptosis-inducing factors, causing cell death.[21] DEX attenuates apoptosis by inhibiting the activation of intrinsic apoptotic cascades.[22]

Nitric oxide synthases (NOS), a family of enzymes catalyzing the production of NO, contribute to cellular damage during IR injury. Different types of NOS differently affect cell viability. NO produced by endothelial NOS (eNOS) has antioxidant properties as it reduces superoxide anion formation, whereas up-regulation of inducible NOS (iNOS) is associated with oxidative stress cytotoxicity. Increased iNOS activity is dependent on activation of ERK-mediated phosphorylation.[23] Here, pre-treatment and post-reperfusion DEX treatment retained phospho-eNOS and decreased phospho-iNOS expression compared to expression in the D-IRC group.

In diabetes, hyperglycemia can activate ROS and the NLRP3 inflammasome. During IR, ROS promote tissue inflammation and activate immune responses through different pathways, including NLRP3 inflammasome.[24] Previously, TXNIP-mediated NLRP3 activation via oxidative stress was identified as a key signaling mechanism in susceptibility to AKI.[16,25] DEX administered to diabetic rats ameliorates lipid peroxidation, oxidative stress, and IR-related renal injury partly by inhibiting P38-MAPK (mitogen-activated protein kinases) activation and expression of TXNIP in diabetic kidney.[7,26] In this study, DEX treatment, especially post-treatment, attenuated increase in oxidative stress markers (CTP1A, NOX4, and TXNIP) following IR.

Furthermore, increased inflammatory markers in IR injury, as expected, were diminished by DEX. In diabetes, elevated levels of circulatory pro-inflammatory factor IL-1β can induce sustained inflammation, causing kidney injury. Reperfusion of ischemic tissues increases inflammatory reactions due to an increase in the release of free radicals and accumulation of inflammatory mediators. Activation of inflammasome also produces IL-1β, a proinflammatory cytokine. DEX was reported to decrease production of cytokines, including IL-1β and TNF-α, in renal IR injury model.[27]

In this study, DEX treatment after IR injury improved renal protection compared to preemptive administration. Therefore, this study provides evidence of retained renoprotective efficacy of DEX under diabetic conditions against IR injury and demonstrates a large therapeutic window of these effects relative to the onset of ischemic insult. Optimal timing of DEX administration remains controversial.[4,28] Gonullu et al. showed that DEX administered before ischemia and after reperfusion histomorphologically reduced renal IR injury, with administration of DEX during reperfusion considered more effective.[29] We speculated that because of the fast onset of effects and rapid clearing of DEX from circulation[30], post-reperfusion treatment with DEX would be more effective in protecting against IR injury than DEX pre-treatment.

No clear recommendations regarding effective DEX dose ranges in experimental IR injury in rat models have been established.[7] However, a wide range of DEX intraperitoneal doses (10-100 μg/kg) showed protective effects against ischemia or IR injury.[6,29,31]

## Conclusions

This study showed that protective effects of DEX in renal IR injury are preserved under diabetic conditions and related to the inflammasome. Furthermore, post-reperfusion DEX treatment was more effective than pre-ischemia treatment. These findings potentially provide a basis for using DEX in treating renal IR injury.

## Acknowledgements

none

## Author contributions

**Conceptualization:** Seung Hyun Kim, Ji Hae Jun, Yong Seon Choi

**Data curation:** Ji Hae Jun, Ju Eun Oh, Eun-Jung Shin, Yong Seon Choi

**Formal analysis:** Seung Hyun Kim, Ji Hae Jun, Ju Eun Oh, Eun-Jung Shin, Yong Seon Choi

**Funding acquisition:** Yong Seon Choi

**Investigation:** Seung Hyun Kim, Ji Hae Jun, Ju Eun Oh, Eun-Jung Shin, Young Jun Oh, Yong Seon Choi

**Methodology:** Seung Hyun Kim, Ji Hae Jun, Yong Seon Choi

**Supervision:** Young Jun Oh, Yong Seon Choi

**Validation:** Seung Hyun Kim, Ji Hae Jun, Young Jun Oh, Yong Seon Choi

**Writing-original draft:** Seung Hyun Kim, Ji Hae Jun

**Writing-review&editing:** Seung Hyun Kim, Ji Hae Jun, Young Jun Oh, Yong Seon Choi

